# Effects of head-only exposure to 900 MHz GSM electromagnetic fields in rats : changes in neuronal activity as revealed by c-Fos imaging without concomitant cognitive impairments

**DOI:** 10.1101/2024.02.20.581017

**Authors:** Bruno Bontempi, Philippe Lévêque, Diane Dubreuil, Thérèse M. Jay, Jean-Marc Edeline

**Affiliations:** Laboratoire de Neurosciences Cognitives, CNRS UMR 5106, Avenue des Facultés, 33405 Talence, France; XLIM, CNRS UMR 6172, Université de Limoges, 123 Av Albert Thomas, 87060 Limoges Cedex, France; UMR CNRS 8195, Centre de Neurosciences Paris-Sud; Université Paris Sud, Batiment 446, 91405 Orsay Cedex, France; INSERM, UMR 894 (ex796), Physiopathologie des Maladies Psychiatriques, Paris 75014, France; Université Paris Descartes, Faculté de Médecine, Paris 75014, France

**Keywords:** Spatial learning, immunohistochemistry, rat, Radio Frequency

## Abstract

Over the last decade, animal models have been used to evaluate the physiological and cognitive effects of mobile phone exposures. Here, we used a head-only exposure system in rats to determine whether exposure to 900MHz GSM electromagnetic fields (EMF) induces regional changes in neuronal activation as revealed by c-Fos imaging. In a first study, rats were exposed for 2h to brain average specific absorption rates (BASARs) ranging from 0.5 to 6W/kg. Changes in neuronal activation were found to be dose-dependent with significant increases in c-Fos expression occurring at BASAR of 1W/kg in prelimbic, infralimbic, frontal and cingulate cortices. In a second study, animals were submitted to either a spatial working memory (WM) task in a radial maze or a spatial reference memory (RM) task in an open field arena. Exposures (45min) were conducted before each training session (BASARs of 1 and 3.5W/kg). Control groups included sham-exposed and control cage animals. In both tasks, behavioral performance evolved similarly in the four groups over testing days. However, c-Fos staining was significantly reduced in cortical areas (prelimbic, infralimbic, frontal, cingulate and visual cortices) and in hippocampus of animals engaged in the WM task (BASARs of 1 and 3.5W/kg). In the RM task, EMF exposure-induced decreases were limited to temporal and visual cortices (BASAR of 1W/kg). These results demonstrate that both acute and subchronic exposures to 900MHz EMFs can produce biological effects, but these effects were not sufficient to induce detectable cognitive deficits in the tasks used here.

## 1. Introduction

Over the last decade, the rapid expansion of mobile communication has raised concerns about the possible deleterious health effects of electromagnetic fields (EMFs) generated both by relay stations and cellular phones (Hermann & Hossmann, 1997; Ahlbom et al., 2001; Hossmann & Hermann, 2003). Although the biological effects of EMFs remain controversial, the brain is likely to be a primary target of EMFs because cellular phones are usually held close to the head while in use. A large diversity of effects has been reported following exposure to various energy levels and types of EMFs which, sometimes, are quite different from those used in global systems for mobile (GSM) communication. In humans, exposure to either low or high frequency EMFs has been suspected to increase the risks of cancer (Caplan et al., 2000; Hardell et al., 2002; Richter et al., 2002) and of leukemia (Juutilainen et al., 1990; Rothman et al., 1996). Impairments of cognitive functions (Preece et al., 1999, Koivisto et al., 2000), changes in electroencephalogram (Croft et al., 2002; Krause et al., 2002a,b; Borbely et al., 1999; Huber et al., 2000, 2002) as well as higher occurrence of headaches (Frey, 1998; Hocking & Westerman, 2001) have also been reported. In animals, where EMF exposure conditions are more controlled, some biological effects have also been documented. Changes in electrophysiological activity (Beason & Semm, 2002), in neurotransmitter content (Lai et al., 1991, 1992, 1993; Chance et al., 1995; Mausset et al., 2001), and in glial reactivity (Mausset-Bonnefont et al., 2004; Brillaud et al., 2007; Ammari et al., 2008a) have been described. However, other studies did not report significant effects: brief or prolonged exposures to EMF were shown to have negligible effect on the blood brain barrier integrity (Fritze et al., 1997; Finnie et al., 2002) and on the genomic response of brain cells (Fritze et al., 1997; Watilliaux et al 2011).

A lack of consensus is also striking in the field of cognitive neuroscience: Some studies have shown an impairment of learning and memory processes in humans (Koivisto et al., 2000) and animals (Lai, 1996; Lai et al., 1994, 1998), whereas others failed in reproducing significant deleterious effects (Davis et al., 1984; Hong et al., 1988; Sienkiewicz et al., 1996, 2000; Dubreuil et al., 2002, 2003; Cassel et al., 2004; Cobb et al., 2004). Several factors could potentially explain these discrepancies, in particular the EMF frequencies and intensities but also the exposure conditions. Indeed, testing rats with the same behavioral tasks, some studies have used whole-body exposures (Lai, 1996; Lai et al., 1994, 1998) whereas others have selected head-only exposures (e.g. see Dubreuil et al., 2002, 2003), this latter situation mimicking better the exposure to cellular phones in human.

The aim of this study was to determine if acute or subchronic exposure to GSM EMFs could induce detectable changes in neuronal genomic response by measuring changes in the expression of the immediate early gene c-Fos. This gene is widely used as a marker of neuronal activity and is also involved in the control of target genes playing a crucial role in the long-term reorganization of brain circuits during various pathogenic and non-pathogenic situations. Using c-Fos imaging, our goal was to identify the different brain regions which may be the most sensitive to EMF exposures.

In a first experiment, we describe the consequences of a single exposure, performed during resting conditions, for groups of rats submitted to brain averaged specific absorption rate (BASAR) ranging from 0.5 to 6 W/kg (i.e, from below the international limit of 2W/kg up to 3 times this value). In a second experiment, we describe the consequence of a subchronic exposure (14 days), performed before behavioral tests, on groups of rats submitted to two BASARs (1 and 3.5W/kg). The rational was to evaluate if repeated exposures lead either to larger effects than a single exposure (by cumulating effects occurring after single exposure) or to smaller effects than a single exposure (by adaptation to EMF exposure). In addition, by engaging the animals in a learning task, one can potentially look for both increases and decreases c-Fos labelling whereas only increases are usually reported in resting conditions.

## 2. Materials and Methods

### 2.1. Animals

Naïve male Sprague Dawley rats (150-180 g at their arrival in the laboratory, Charles River, L’Arbresle, France) were housed in a humidity (50-55%) and temperature (22-24° C)-controlled facility on a 12 h:12 h light/dark cycle (lights on at 6:30 A.M.) with free access to food and water, except during behavioral testing. Animals were group-housed (two to five per cage) and were allowed a 1-week period of habituation to the colony room before testing. They were handled and weighed every day. At the end of the habituation period, rats were single housed. The analysis of the effects of acute EMF exposure (Experiment 1) was conducted in animals that had *ad libitum* access to food and water. Animals used for memory testing (Experiment 2) were progressively food deprived and maintained to 85% of their free-feeding weight. All procedures were performed in conformity with French (JO 887-848) and European (86/609/EEC) legislations on animal experimentation.

### 2.2. Electromagnetic field exposure system

The exposure system was the one used in previous experiments (Dubreuil et al., 2002; 2003); it was described in details in Lévêque et al. (2004). Briefly, a radiofrequency generator (RFPA S.A., model RFS9001800-25), emitting a GSM EMF (1/8 duty factor) pulsed at 217 Hz (pulse emission during 576 sec every 4.6 ms) was connected to a 4-output divider, allowing exposure of four animals simultaneously. Each output was connected to a loop antenna enabling a local exposure of the animal’s head. The loop antenna consisted of a printed circuit, with two metallic lines engraved in a dielectric epoxy resin substrate (dielectric constant ε_r_ = 4.6). At one end, this device consisted of a 1 mm-width line forming a loop placed close to the animal’s head.

Brain average specific absorption rates (BASARs) were determined numerically using a numerical rat model with Finite Difference Time Domain method and experimentally in homogeneous rat phantom using a Vitek or Luxtron probe for the measurement of temperature rises. In this case, BASARs, expressed in W/kg, were calculated using the following relation: SAR = CΔT/Δt, with C being the calorific capacity in J/(kg.K), ΔT, the temperature variation in °K and Δt, the time variation in second. Comparisons were done between numerical and experimental SAR values in homogeneous phantoms, especially in the equivalent rat brain area. The agreement among numerical and experimental SAR values was good (see for details, Lévêque et al., 2004). Finally, the brain average SAR (BASAR) in non-homogeneous rat model was obtained: 6.8W/kg for one Watt incident power. The incident power was than adjusted to obtain the experimental BASAR levels used in our experiments.

EMF exposure and behavioral testing were performed in two different rooms. During radiofrequency exposure, rats were restrained in rockets made of 5 mm-thick Plexiglas and consisting of a cylinder (6 cm in diameter, 15 cm long) and a truncated cone (3 cm long) in which the rat’s head was inserted (see for details Dubreuil et al., 2002, 2003, Lévêque et al. 2004). The loop antenna was placed against the cone directly above the rat’s head. The end of the cone was opened and holes were made in the rocket’s cylinder to facilitate breathing and minimize sweating and increases in body temperature. A Plexiglas disk was secured at the entry of the cylinder to prevent the rat from backing out of the rocket and to minimize as much as possible any movement of the head. To prevent interference from one antenna to another, rockets were separated from each other by a double layer of absorbing material (APM 12, Hydral, 12 cm-thick).

### 2.3. Habituation to restrained conditions

To minimize the nonspecific effects of stress induced by the rocket confinement on learning performance as well as on c-Fos expression, animals were first submitted to a habituation protocol. Rats were placed in the rockets with the loop antenna disconnected from the radiofrequency generator. They were adapted gradually to the restrained conditions for either 14 days (Experiment 1) or 7 days (Experiment 2). Also, on the first day after arrival in the laboratory, rockets were introduced in the home cages, allowing animals to enter freely in them. During the following days, confinement in the rockets was progressively increased from 5 min to 2 hours (Experiment 1) or from 5 min to 45 min (Experiment 2). Across days, the animal movements in the rockets were less frequent attesting that the restrained conditions were relatively well tolerated. In a few cases (n=3), animals showing important movements in the rocket at the end of this habituation period were discarded from the study.

### 2.4. Experiment 1

This experiment was designed to investigate the effects of 900MHz EMF exposure on the basal level of neuronal activity. Awake, “resting” animals (maintained in their home cage prior to being placed in the rockets) were used. Following the 2-weeks habituation to the rockets, animals were randomly assigned to 6 different treatment groups (n = 9/group): one sham-exposed group immobilized for 2 hours in the rockets but without EMF and five experimental groups exposed for 2 hours to 900 MHz EMF at BASARs of 0.5, 1, 2, 4 and 6 W/kg respectively. This range of BASARs was selected because it started below the international limite up to 3 times this value (the SAR limit is set by international requirements and by the Council of the European Union at 2 watts/kilogram averaged over 10 grams of body tissue). The exposure duration was set at 2h which is probably the longer time a rat can accept restrained conditions in the rocket used for the exposure. It is also close to the average time per day for extensive cell phone users.

Immediately at the end of the exposure period, the animals were deeply anesthetized, perfused and their brains were processed for immunocytochemistry as described below.

### 2.5. Experiment 2

#### 2.5.1. Aim and groups

This experiment was designed to determine the effects of EMF exposures on c-Fos expression while rats were submitted to learning tasks. Animals were engaged in behavioral tasks to allow for (i) detection of potential decrease in neuronal activity that will be difficult to observe in basal state and (ii) possible changes in the activation of brain areas in response to cognitive demand. Animals were randomly assigned to four different treatment groups (n=12/group). Cage control rats were maintained in their home cage and did not receive any treatment before being submitted to the learning task. Sham exposed rats were immobilized for 45 min in the rockets immediately prior to each daily training session but without EMF exposure. Two experimental groups were exposed to 900 MHz EMF at BASAR of either 1 or 3.5 W/kg for 45 min immediately prior to each daily training session. The BASAR of 1W/kg was selected based on the results of experiment 1 and the BASAR of 3.5 W/kg was the higher our system can delivered when 4 animals were simultaneously exposed^1^. There was a time-lag of 20 minutes between the beginning of exposure of the four animals (i.e., they were placed in the rockets every 20 min) thus allowing to test each rat immediately at the end of the EMF exposure or the sham exposure. Time of testing in the day (morning or afternoon) was counterbalanced between groups. The equipment and the behavioral procedures used in the two tasks were exactly the same as those described in our previous experiments Dubreuil et al. (2002, 2003b). They are only briefly summarized here.

#### 2.5.2. Spatial working memory task

This task was conducted in a radial maze made of an octagonal central platform (23 cm in diameter) from which radiated eight arms (80 cm long, 12 cm wide) in a symmetrical fashion. At the end of each arm, a food cup (5 cm in diameter, 1 cm in height) could be baited with food rewards (i.e. chocolate cereals). The room was illuminated by a neon light located above the apparatus, thus avoiding shadows within the maze. Large cues were placed on the walls of the room and serve as spatial cues. In order to be familiarized with the radial maze and its environment, rats were first submitted to habituation sessions immediately after the last 3 days of adaptation to restrained conditions. After the restraint conditions in the rockets, a group of six rats was placed for 20 minutes in the maze where food rewards were scattered throughout the maze to encourage exploration. On the last day of habituation, rats explored the maze in groups of three and food rewards were placed only in the food cup of each arm. This habituation session was terminated when all eight arms were visited and all food rewards were consumed. Spatial working memory testing sessions began the next day and lasted for 10 consecutive days. Rats were submitted to daily session during which all arms were baited. Each session consisted of one trial and started by placing the rat in an opaque plexiglas cylinder located in the centre of the maze platform. After 10s, the cylinder was removed and the rat was allowed to freely explore the maze until all eight arms had been visited or when the rats had made a total of 16 visits or when 10 min had elapsed. Arms were not rebaited during the ongoing trial, so repeated entries into an arm were counted as working memory errors. Choice accuracy was also evaluated by the rank of the first error which was defined as the number of consecutive correct visits in a training session before the first error occurred (i.e. the maximum rank was 8). Rats learning this task typically show a progressive decrease in the number of working memory errors over days while the rank of the first error increases. Seventy to ninety minutes after the end of the 10^th^ session, half of the animals randomly selected (n=6/groups) were deeply anesthetized, perfused and the brains were processed for immunocytochemistry (see below).

#### 2.5.3. Spatial reference memory task

Reference memory performance was measured using a dry-land version of the Morris water maze (Lavenex & Schenk, 1995; Delatour & Gisquet-Verrier 2000, Dubreuil et al., 2002). This spatial navigation task was conducted using a circular arena (120 cm in diameter) surrounded by a transparent plastic edge (10 cm high). The arena was mounted on a rotating pedestal, 70 cm above the floor. Thirteen white opaque circular boxes (3 cm in diameter) that were glued on the arena, defined a regular geometrical pattern in the arena. Each box was covered by a white lid. Chocolate cereals used as food rewards were placed in selected boxes. During the last 4 days of adaptation to restrained conditions in the rockets, rats were familiarized with the boxes placed in the arena. On the first day, rats were habituated to eat the food reward contained in a box placed in their home cage. On the two following days, the box was covered with the lid and the rats had to learn to open it to get the food reward. Rats were given four trials/day for two consecutive days. On the second day, rats had to open the box in less than 10 s otherwise four additional trials were given. On the fourth habituation day, rats were placed by group of three on the circular platform and were free to explore the non-baited apparatus for 15 min. Spatial reference memory testing session begun the next day and lasted for 14 consecutive days. Each rat had to learn that the food reward was located in only one box. For a given animal, the baited box was always located at the same position with regard to the reference space formed by the geometrical cues scattered around the room. Rats were submitted to a daily session made of four consecutive trials separated by a 1-min intertrial interval. The rat was placed on the platform at each of four different starting positions (North, South, East and West) facing the transparent wall of the platform. From day to day, the order of the starting positions was randomly changed to prevent the rats from learning a specific route. To prevent strategies based on olfactory cues, between each trail and each rat, the platform was randomly turned by 60, 120, 180, 240 and 300°. Each trial lasted a maximum of 5 min. If the rat did not find the goal box after 5 min, it was gently guided in front of the goal box and allowed to open it and eat the food reinforcement. Reference memory errors were defined as the number of boxes, other than the goal box, that the rat opened during a trial. A hit was achieved when the rat opened the goal box directly without opening any other box; on a given day the maximum number of hits was four. Latency to open the goal box was also measured. Seventy to ninety minutes after the end of the 14^th^ session, half of the animals randomly selected (n=6/groups) were deeply anesthetized, perfused and the brains were processed for immunocytochemistry (see below).

### 2.6. c-Fos immunocytochemistry

Following deep anesthesia, rats were perfused transcardially with 350 ml of 0.9% NaCl followed by 350 ml of cold 4% paraformaldehyde in 0.1 M phosphate buffer (PB), pH 7.4. Brains were dissected, post-fixed for 12 hours in the same fixative and cryoprotected in 30% sucrose/PB and left overnight at 4°C. They were then frozen and 50 µm frontal sections were cut on a freezing microtome. Immunohistochemistry was performed on free-floating sections using a standard avidin-biotin (ABC Standard Elite kit, Vector Laboratories, Burlingame, CA, USA) method. The primary antibody was a rabbit polyclonal antibody (Ab-5, Oncogene Science, 1:20000) raised to a synthetic peptide derived from aminoacid sequences 4-17 of the Fos protein. Incubation in 2% goat serum containing 0.2% Triton X-100 (Sigma) and the primary antibody occurred overnight at room temperature on a rotating shaker. Following a series of 4 washes in PB, sections were incubated for 2 h at room temperature in a solution containing a biotinylated goat anti-rabbit secondary antibody (Jackson Immunoresearch, 1:2000). Sections were washed again and processed with the avidin-biotin complex for 2 hr at room temperature. Diaminobenzidine (0.05 % solution, Sigma) was used as chromogen and revealed using a few drops of fresh 0.3% hydrogen peroxide solution. The reaction was stopped by washing in PB. Sections were mounted on gelatin-coated slides, dehydrated through a series of graded alcohols baths and coverslipped.

### 2.7. Image analysis and cell counting

Quantitative analysis of c-Fos-positive nuclei was performed using a color video camera (Sony® DXC-950P) interfaced with an Olympus® BX 50 microscope. Fos-positive nuclei were counted in selected brain regions using computer-assisted software (Biocom Visiol@b® 2000; x20 magnification). Sampled areas defined according to the atlas of Paxinos and Watson (1986) are presented in Figure 1. The number of c-Fos-positive nuclei measured in each region was counted bilaterally using a minimum of three consecutive sections in each brain by experimenters who were blind to the exposure conditions. The mean numbers of c-Fos-positive nuclei contributed by each animal in a group were averaged to generate the final means. Data are expressed as the number of c-Fos-positive nuclei/mm^2^ for all regions studied.

**Figure 1.**
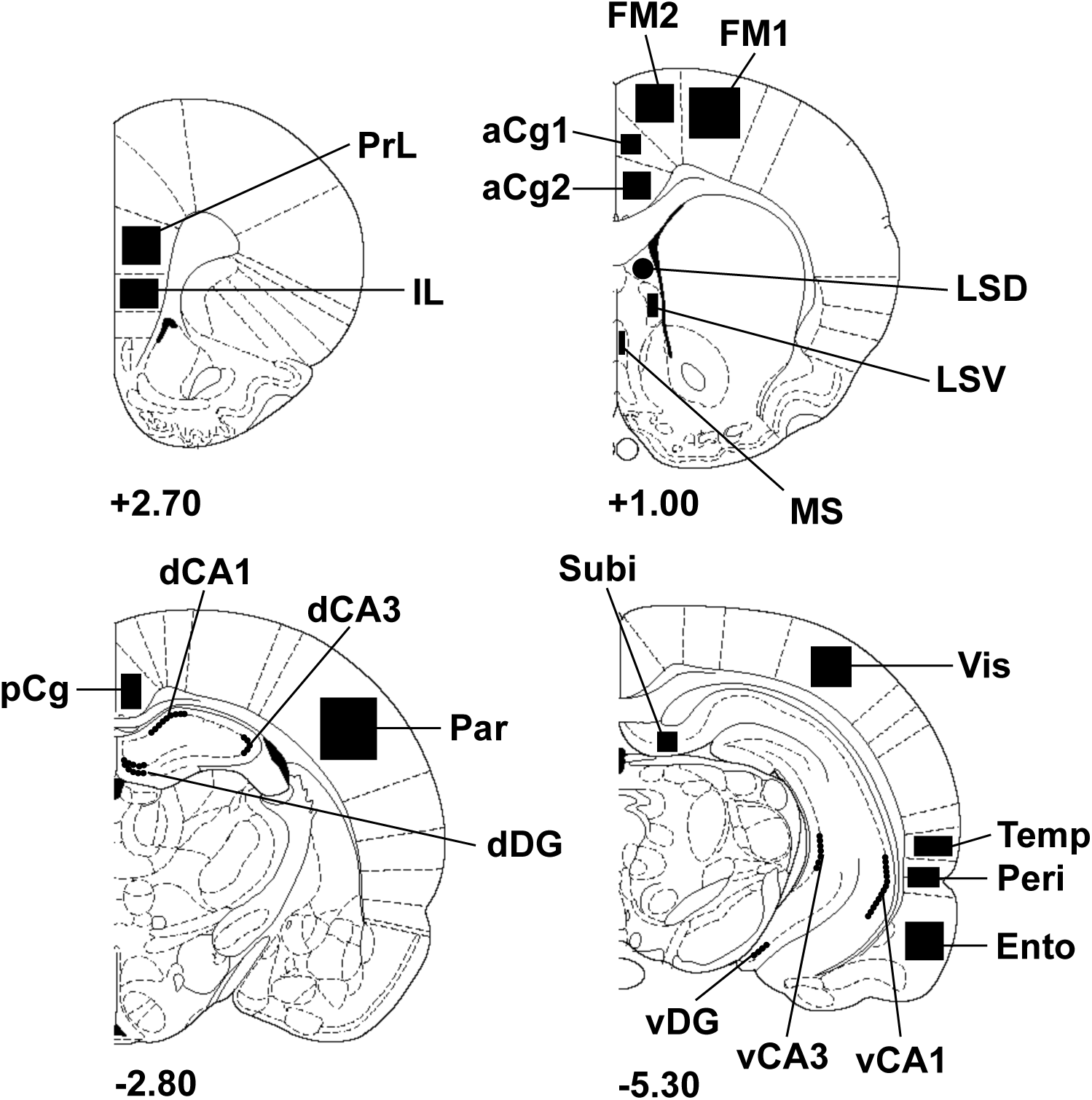
Schematic drawings of rat brain coronal sections showing the regions of interest (filled areas) selected for c-Fos imaging (adapted from the atlas of Paxinos and Watson,1986). Numbers indicate the distance in millimeters of the sections from bregma. aCg1: anterior cingulate cortex, area 1; aCg2: anterior cingulate cortex, area 2; dCA1: dorsal hippocampus, CA1 field; dCA3: dorsal hippocampus, CA3 field; dDG: dorsal dentate gyrus; Ento: entorhinal cortex; FM1: frontal cortex, primary motor area 1; FM2: frontal cortex, secondary motor area 2; IL, infralimbic cortex; LSD: lateral septum, dorsal part; LSV: lateral septum, ventral part; MS: medial septum; Par: parietal cortex; pCg: posterior cingulate cortex; Peri: perirhinal cortex; PrL, prelimbic cortex; Subi: subiculum; Temp: temporal associative cortex; vCA1: ventral hippocampus, CA1 field; vCA3: ventral hippocampus, CA3 field; vDG: ventral dentate gyrus; Vis: visual cortex, secondary lateral area.

### 2.8. Statistical analyses

Behavioral parameters and numbers of c-Fos-positive nuclei were compared using an analysis of variance (ANOVA) with repeated measures and, when appropriate, were followed by *post hoc* paired comparisons using the Newman-Keuls test. All data are presented as mean ± SEM. For all comparisons, values of p < 0.05 are considered as statistically significant.

## 3. Results

### 3.1. Experiment 1: Effect of EMF exposure on neuronal activity as revealed by c-Fos imaging

A two-way ANOVA performed on c-Fos mapping data with BASARs as between group factor and the 23 brain regions analyzed as within-subjects factor indicated a significant group x region interaction (F(5, 1190) = 7.11; p < 0.0001), showing that region-specific changes in c-Fos expression occurred following EMF exposure (see Figure 2). Analysis of individual regions revealed that EMF exposure induced a dose-dependent increase in c-Fos expression in the following brain regions: subiculum (F(5, 46) = 2.47; p < 0.05), prefrontal (prelimbic area, F(5,46) = 2.49; p < 0.05; infralimbic area, F(5, 46) = 3.78; p < 0.01), frontal (M2 motor area, F(5, 46) = 2.73; p < 0.05) and anterior cingulate cortices (area 2, F(5, 46) = 2.51; p < 0.05).

**Figure 2.**
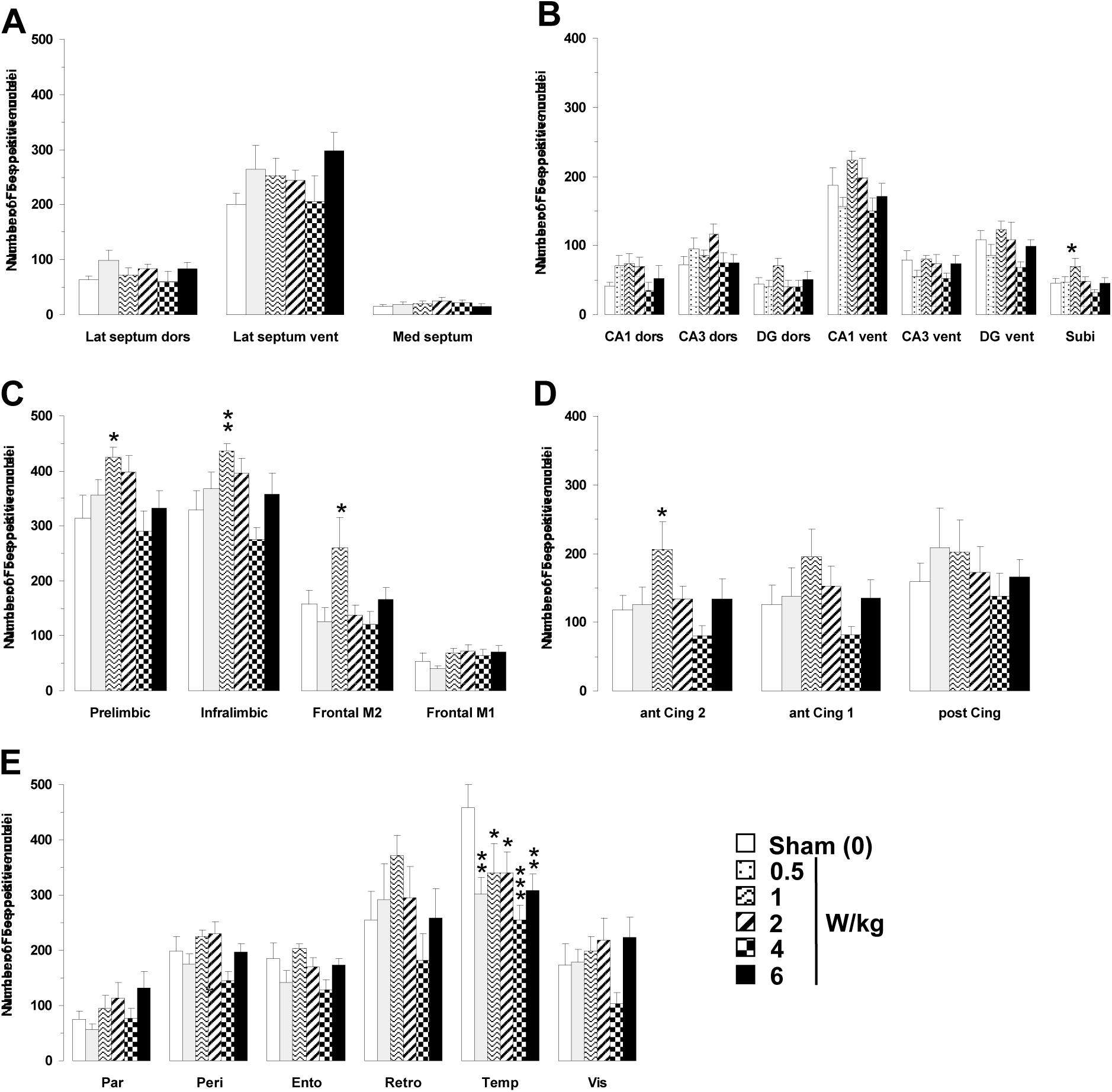
Effects of acute GSM 900 MHz electromagnetic field exposure on neuronal activity as measured in 23 brain areas by c-Fos immunohistochemistry in exposed and sham-exposed animals. Data are presented as number of c-Fos-positive nuclei in selected brain regions (mean ± SEM, n = 9/group). A, septal region; B, hippocampal formation, C, D, E, cortical regions. See Figure 1 for list of abbreviations. Experimental animals were exposed for 2 h while sham-exposed animals were placed in rockets but without any EMF exposure. *p<0.05, **p<0.01, ***p<0.001 for exposed versus sham-exposed animals, ANOVA followed by Fisher post hoc test.

*Post-hoc* analyses revealed that only the BASAR of 1 W/kg significantly increased c-Fos expression as compared to sham animals in the subiculum (+ 53.2%, p < 0.05), prefrontal (prelimbic area + 35.2%, p < 0.05; infralimbic area + 32.9%, p < 0.01), frontal (M2 motor area + 64.9%, p < 0.05) and anterior cingulate cortices (area 2, + 74.2%, p < 0.05; see Figure 2). A reduction in c-Fos expression was found in the temporal cortex at all BASARs (F(5, 46) = 3.26; < 0.05), with the greatest decrease observed at the BASAR of 4 W/kg (-44.4%, p < 0.001; Figure 2). The perirhinal cortex was similarly affected (F(5, 46) = 2.87; p < 0.05) with a significant decrease observed at the BASAR of 4 W/kg (-27.3%, p < 0.05). Examples of the largest EMF-induced increase in c-Fos expression in the prefrontal and frontal cortices are presented in Figure 3.

**Figure 3.**
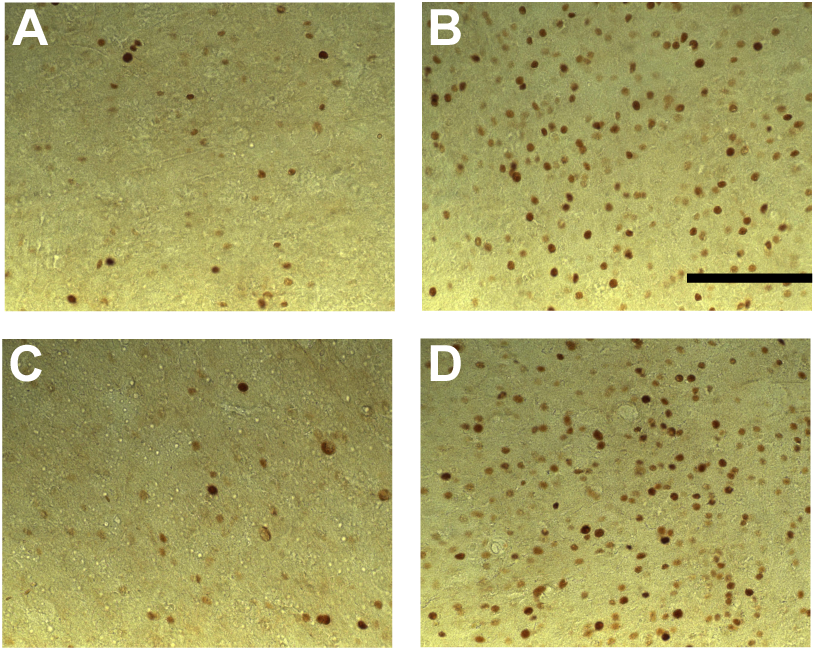
Photomicrographs of c-Fos immunoreactivity in coronal sections taken through the prelimbic cortex (A, B) and frontal cortex (area FM2; C, D) following acute GSM 900 MHz electromagnetic field exposure (SAR of 1 W/kg for 2 hr). Note that this example represents the largest increased number of c-Fos-positive nuclei in exposed animals (1 W/kg; B, D) as compared to sham-exposed animals (0 W/kg; A, C). Scale bar: 100 µm.

### 3.2. Experiment 2: Effect of EMF exposure on learning and memory performance and associated neuronal activity

#### 3.2.1. Behavioral Data

In the following paragraphs, we present the behavioral results obtained from the subset of animals (n=6 per group) used for the quantification of c-Fos labeling. In all but one cases, these results are similar to those obtained with the whole population of animals ran in the behavioral tests (n=12 per group, see supplementary figures 1 and 2).

##### 3.2.1.1. Working memory task

As shown in Figure 4A, EMF exposure did not affect significantly the number of working memory errors recorded over 10 days of testing (Treatment F(3, 20) = 2.06, p > 0.13, NS). For all groups, the number of working memory errors decreased significantly over days (F(9, 180) = 3.08, p < 0.002) and the slope of the learning curve was similar across groups (Treatment x Day interaction: F(27, 180) = 1.06, p > 0.38, NS). As illustrated in Figure 4B, the rank of the first error, which can be taken as a relevant index of working memory performance, was similar across groups (F(3, 20) = 0.74, p > 0.54, NS) and increased in a similar way over days (Day: (F9, 180) = 2.54; p < 0.01); Treatment x Day interaction: F(27, 180) = 1.22; p > 0.22, NS). Latency to complete the task was also similar across groups (data not shown). The same results were obtained when analyses were performed on the entire population of animals used in this study (n=12 per group, see supplementary figure 1).

**Figure 4.**
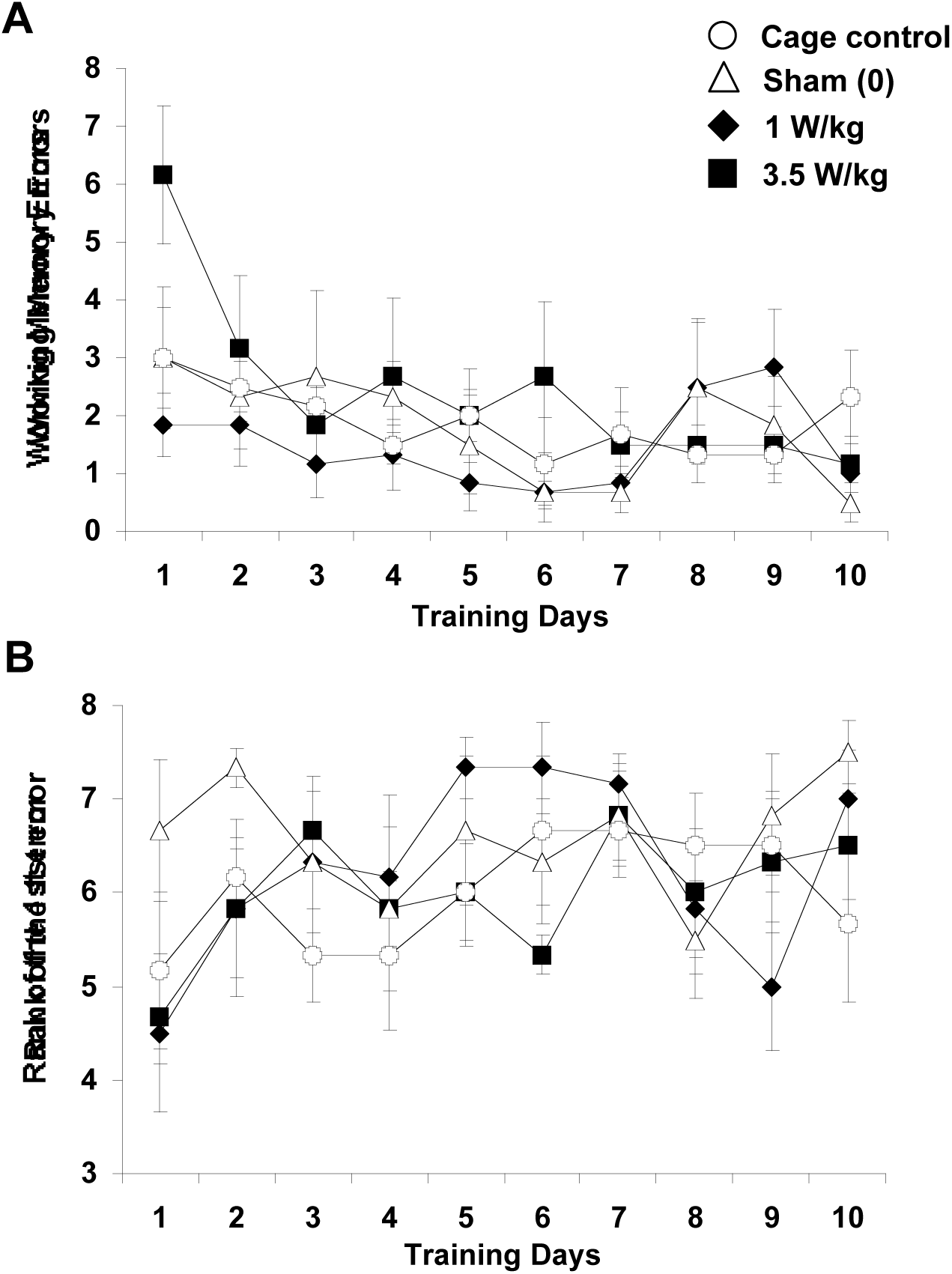
Effect of subchronic GSM 900 MHz electromagnetic field exposure on working memory performance as measured in the 8-arm radial maze. Data are presented as number of working memory errors (A) and rank of the first error (B) (mean ± SEM, n = 6/group) measured over 10 days of training. Cage control rats were not submitted to any treatment before each daily training session. Sham-exposed rats were placed in the rockets for 45 min without any EMF exposure. Experimental animals were exposed to EMFs for 45 min at SARs of either 1 or 3.5 W/kg. Note that for the rank of the 1^st^ error, a high value indicates a better performance. No significant between group differences were observed.

##### 3.2.1.2. Reference memory task

As shown in Figure 5A, EMF exposure did not affect significantly the number of reference memory errors recorded over 14 days of testing (Treatment F(3, 20) = 1.87; p > 0.16, NS). For all groups, the number of errors decreased significantly as training progressed (F13, 260) = 10.17; p < 0.0001) and the slope of the learning curve was similar (Treatment x Day interaction: F(39, 260) <1, NS). The number of hits (Figure 5B), which can be taken as a relevant index of spatial discrimination performance, increased significantly from Day 1 to Day 14 ((F13, 260) = 8.99, p < 0.001) and evolved similarly for all groups (Treatment x Day interaction: F(39, 260) <1, NS). However, there was a main effect of EMF exposure (Treatment: F(3, 20) = 4.07, p < 0.03). Subsequent analyses revealed that, although the 1W/kg exposed group did not differ from the control group (F<1; NS), it significantly differed from the sham exposed group (F(1,10)=3.45; p <0.05). This difference probably comes from the fact that the 1W/kg group exhibited a lower number of hits than the sham group on days 3 and 8 (p<0.05). However, this difference between animals exposed to 1W/kg and sham-exposed animals was not found on the entire population of animals used in this study (n=12 per group; supplementary figure 2). On the whole population, there was no “treatment” effect, nor any interaction between the “day” and treatment factors, when the whole population of animals was considered (Fs <1; NS).

**Figure 5.**
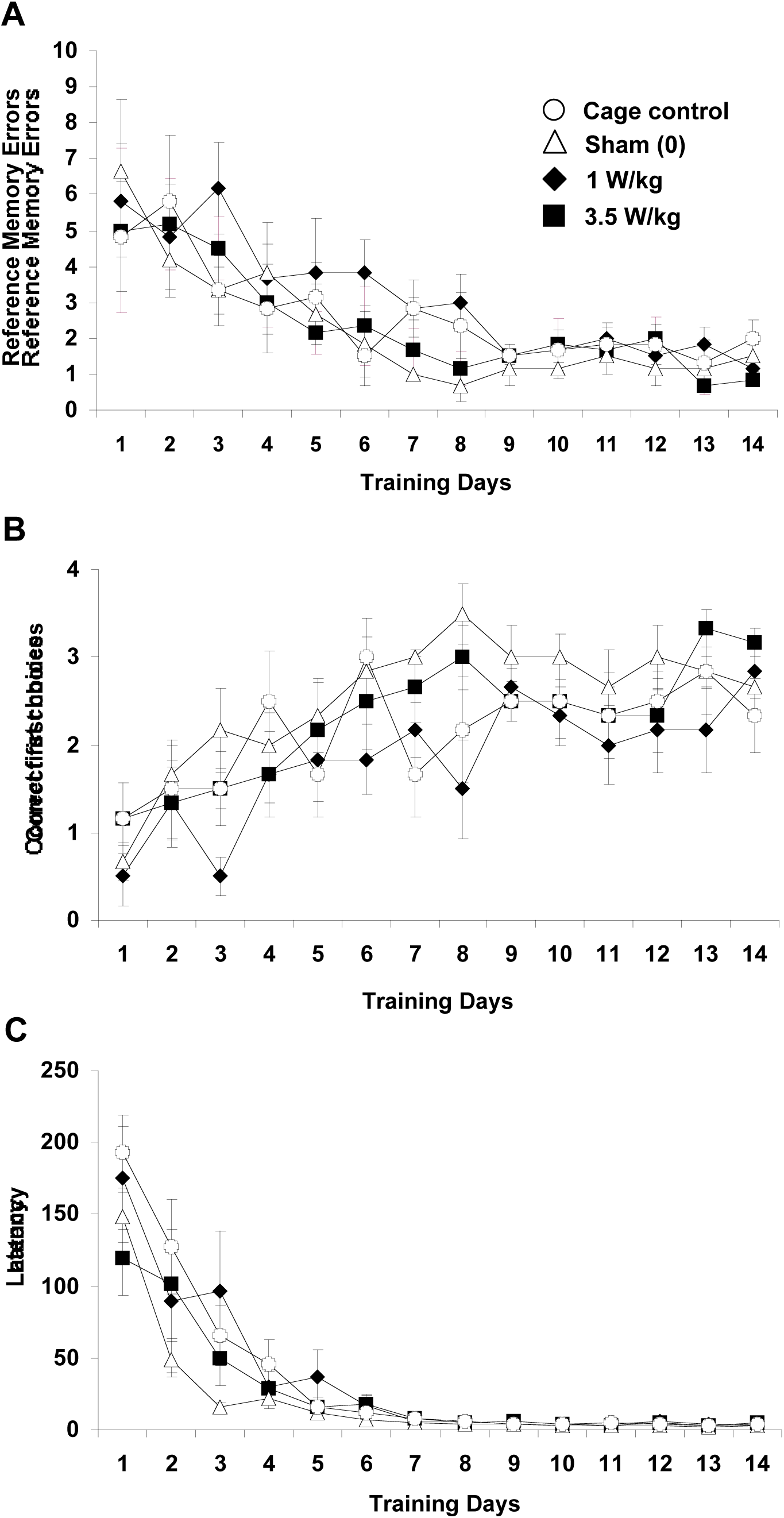
Effect of subchronic GSM 900 MHz electromagnetic field exposure on reference memory performance as measured in the spatial navigation task (dry-land version of the Morris water maze). Data are presented as number of reference memory errors (A), number of hits (B) and latency in sec to find the baited box (C) (mean ± SEM, n = 6/group) measured over 14 days of training. Cage control rats were not submitted to any treatment before each daily training session. Sham-exposed rats were placed in the rockets for 45 min without any EMF exposure immediately prior to each testing session. Experimental animals were exposed to EMFs for 45 min at SARs of either 1 or 3.5 W/kg immediately prior to each testing session. No difference was observed in terms of number of errors (A) and latency to reach the goal box (C). In terms of number of hits (B), the group exposed to 1W/kg differed from the sham exposed group but did not differ from the cage control group. This effect on the number of hits was not found when larger sets of animals (n=12 per group) were tested.

As illustrated in Figure 5C, the latency to find the goal box was also similar across groups (Treatment F(3, 20) = 1.15, p > 0.35, NS) and decreased in a similar fashion over days (Day: (F(13, 260) = 51.41, p < 0.0001); Treatment x Day interaction: F(39, 260) = 1.30; p > 0.12, NS). A more detailed analysis performed on the four trials of a daily session did not show any significant differences between groups, whatever the day of learning considered, thus indicating that learning occurring within a session was similar across groups (data not shown).

#### 3.2.2. c-Fos mapping

##### 3.2.2.1. Working memory task

As a general rule, no significant differences in c-Fos expression were observed between cage control animals and sham animals placed in the rockets without any EMF exposure. As shown in Figure 6, the only increase after sham-exposure was observed in the infralimbic area of the prefrontal cortex (+33.9%; p < 0.01). A two-way ANOVA performed on c-Fos mapping data with BASARs as between group factor and the 23 brain regions analyzed as within-subjects factor indicated a significant group x region interaction (F(3, 500) = 3.87; p < 0.01), showing that region-specific changes in c-Fos expression occurred following EMF exposure. Analysis of individual regions revealed that EMF exposure induced a dose-dependent decrease in c-Fos expression in the following brain regions: hippocampus (CA1 ventral part (F(3, 20) = 3.56; p < 0.05), prefrontal (prelimbic area, F(3, 20) = 3.10; p < 0.05; infralimbic area, F(3, 20) = 6.97; p < 0.05), frontal (M2 motor area, F(3, 20) = 3.34; p < 0.05), anterior cingulate (area 2, F(3, 20) = 3.08; p < 0.05), perirhinal (F(3, 20) = 3.12; p < 0.05) and visual cortices (F(3, 20) = 5.76; p < 0.01).

**Figure 6.**
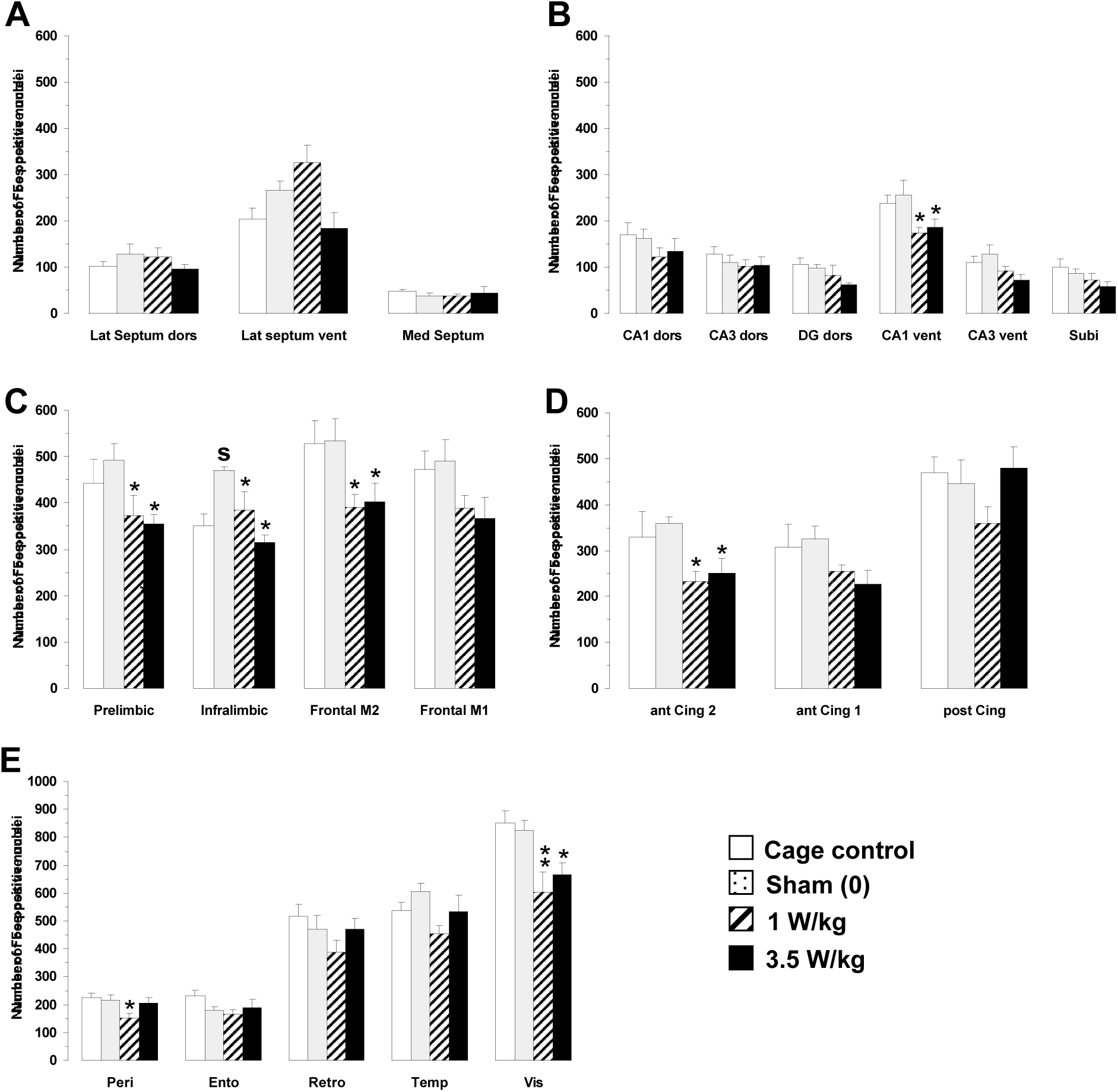
Effects of subchronic GSM 900 MHz electromagnetic field exposure on neuronal activity as measured by c-Fos immunohistochemistry in cage control, exposed and sham-exposed animals submitted to the working memory task. Data are presented as number of c-Fos-positive nuclei in selected brain regions (mean ± SEM, n = 6/group). A, septal region; B, hippocampal formation, C, D, E, cortical regions. See Figure 1 for complete list of abbreviations. Cage control rats were not submitted to any treatment before each daily training session. Sham-exposed rats were placed in the rockets for 45 min without any EMF exposure immediately prior to each testing session. Experimental animals were exposed to EMFs for 45 min at SARs of either 1 or 3.5 W/kg immediately prior to each testing session. *p<0.05, **p<0.01, ***p<0.001 for exposed versus sham-exposed animals. ^S^p < 0.05 for sham-exposed versus cage control animals, ANOVA followed by Fisher post hoc test.

Post-hoc analyses revealed that EMF exposure significantly decreased c-Fos expression as compared to sham animals in the following regions (Figure 6): hippocampus (CA1 ventral part, BASAR of 1 W/kg - 32.1%, p < 0.05; SAR of 3.5 W/kg – 26.9%, p < 0.05), prefrontal (prelimbic area, BASAR of 1 W/kg - 24.4%, p < 0.05; BASAR of 3.5 W/kg – 27.9%, p < 0.05; infralimbic area, BASAR of 1 W/kg - 18.1%, p < 0.05; BASAR of 3.5 W/kg – 32.9%, p < 0.001), frontal (M2 motor area, BASAR of 1 W/kg - 26.9%, p < 0.05; BASAR of 3.5 W/kg – 24.7%, p < 0.05), anterior cingulate (area 2, BASAR of 1 W/kg - 35.3%, p < 0.05; BASAR of 3.5 W/kg – 29.9%, p < 0.05), perirhinal (BASAR of 1 W/kg - 29.3%, p < 0.05), and visual cortices (BASAR of 1 W/kg - 26.8%, p < 0.01; BASAR of 3.5 W/kg – 19.1%, p < 0.05). Examples of the largest EMF-induced decreases in c-Fos expression in frontal cortex are presented in Figure 7.

**Figure 7.**
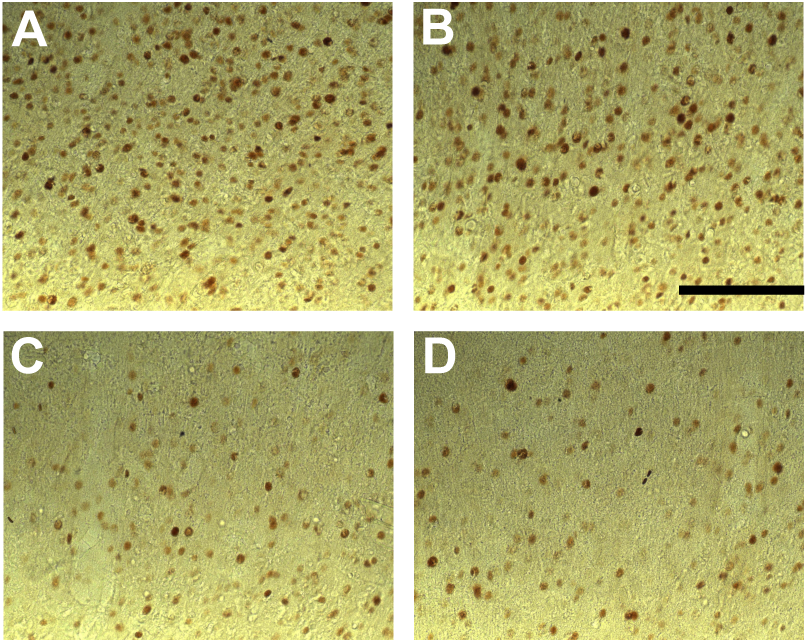
Photomicrographs of c-Fos immunoreactivity in coronal sections taken through the frontal cortex (M2 area) of animals submitted to the working memory task for 10 days and exposed daily to GSM 900 MHz electromagnetic fields (SAR of 1 or 3.5 W/kg). Exposure was given for 45 min immediately prior to each testing session. **Note that this example represents the largest** decreased number of c-Fos-positive nuclei in exposed animals (1 and 3.5 W/kg; C and D, respectively) as compared to cage control (A) and sham-exposed animals (0 W/kg; B). No significant differences were observed between cage control and sham-exposed animals. Scale bar: 100 μm.

##### 3.2.2.2. Reference memory task

As shown in Figure 8, no significant differences in c-Fos expression were observed between cage control animals and sham animal placed in the rockets without exposure to EMFs. As for animals engaged in the working memory task, one-way ANOVA revealed that EMF exposure induced a dose-dependent decrease in c-Fos expression but in only two brain regions: the temporal (F(3, 20) = 3.05; p < 0.05) and visual cortices (F(3, 20) = 3.30; p < 0.05). Post-hoc analyses revealed that exposure to a BASAR of 1 W/kg significantly decreased c-Fos expression in the temporal cortex by 23.6% (p < 0.05) and in the visual cortex by 18.5% (p < 0.05) as compared to sham animals (Figure 8E).

**Figure 8.**
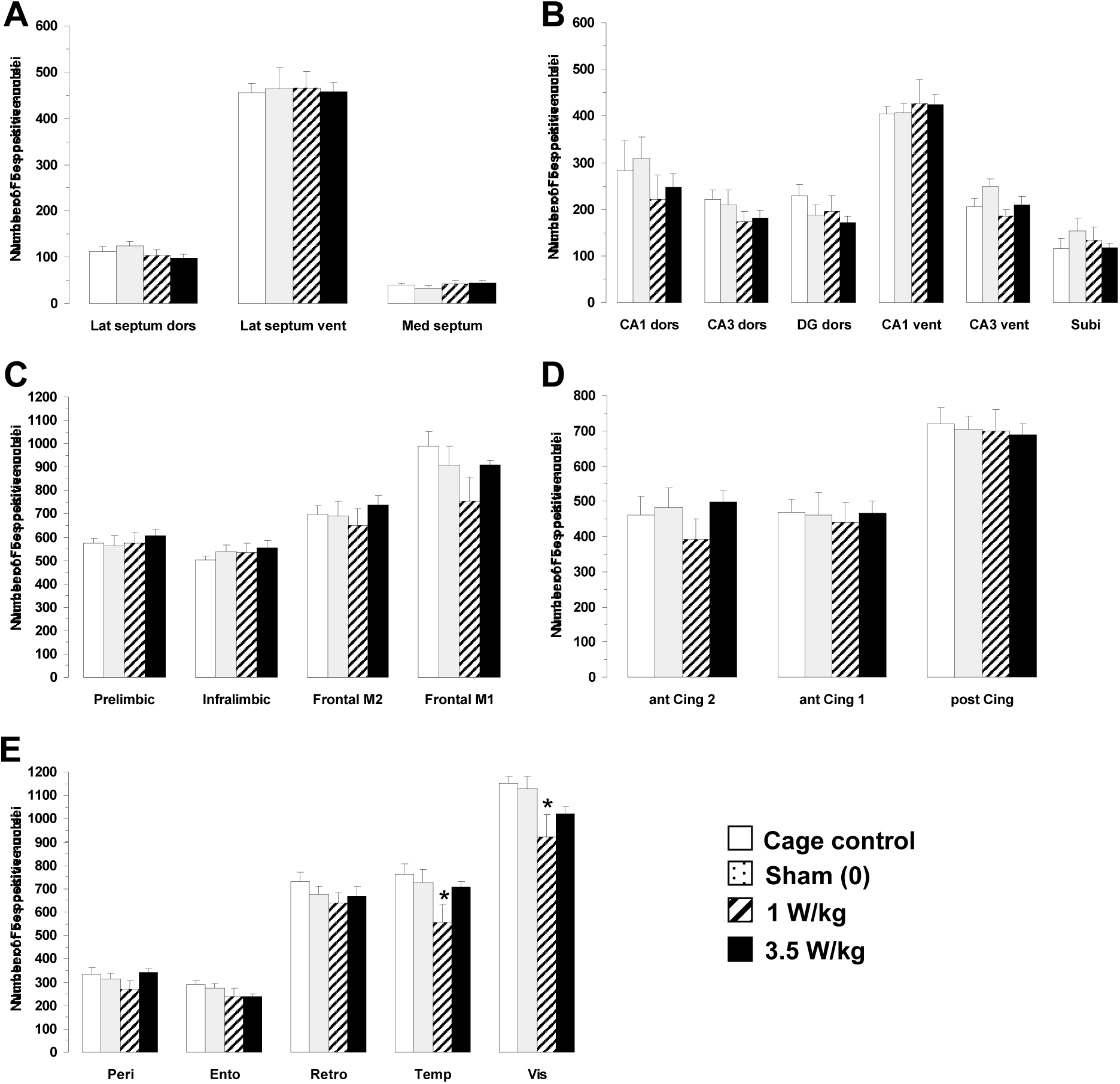
Effects of subchronic GSM 900 MHz electromagnetic field exposure on neuronal activity as measured by c-Fos immunohistochemistry in cage control, exposed and sham-exposed animals submitted to the reference memory task. Data are presented as number of c-Fos-positive nuclei in selected brain regions (mean ± SEM, n = 6/group). A, septal region; B, hippocampal formation, C, D, E, cortical regions. See Figure 1 for complete list of abbreviations. Cage control rats were not submitted to any treatment before each daily training session. Sham-exposed rats were placed in the rockets for 45 min without any EMF exposure immediately prior to each testing session. Experimental animals were exposed to EMFs for 45 min at SARs of either 1 or 3.5 W/kg immediately prior to each testing session. *p<0.05, **p<0.01, ***p<0.001 for exposed versus sham-exposed animals, ANOVA followed by Fisher post hoc test.

### 3.3. Relations between dosimetric analysis and changes in c-Fos expression

In the two experiments presented here (i.e. both after acute and subchronic exposures), some brain areas exhibited changes in c-Fos labeling whereas many others did not. Two possibilities could explain why only a few areas show changes in c-Fos labeling. First, these areas might be those where the local SAR values are the highest. Second, these areas might be those indirectly activated by EMF exposures via mechanisms that remain to be determined.

To search for relationships between the SAR values and the cortical areas exhibiting changes in c-Fos activation after EMF exposure, we referred to SARs levels calculated in a non-homogeneous rat model with a 1-mm^3^ spatial resolution (detailed in Lévêque et al., 2004). Figure 9A and 9B give global views of the rat brain model (with the loop antenna above) and show that the rat’s head absorbs 86% of the total power absorbed by the animal. For eleven brain areas showing changes in c-Fos activation in our experiments (see supplementary Table 1), the Paxinos and Watson (1986) coordinates were converted in coordinates used for the rat brain model and their local SARs were determined. Figure 9C presents the SAR distribution in the rat brain from four sagittal sections and indicates the location of 4 areas (by a black square). It appears that the prelimbic and infralimbic areas (location 1 and 2) receive low SAR values whereas the two frontal areas M2 and M1 (location 3 and 4) receive high SAR values. Yet, these 4 areas exhibited changes in c-Fos labeling during resting conditions (experiment 1) and during behaviorally-induced c-Fos activation (experiment 2). Thus, there was no evidence for systematic relationships between the SAR level in the rat brain and the changes in c-Fos labeling after EMF exposure.

**Figure 9.**
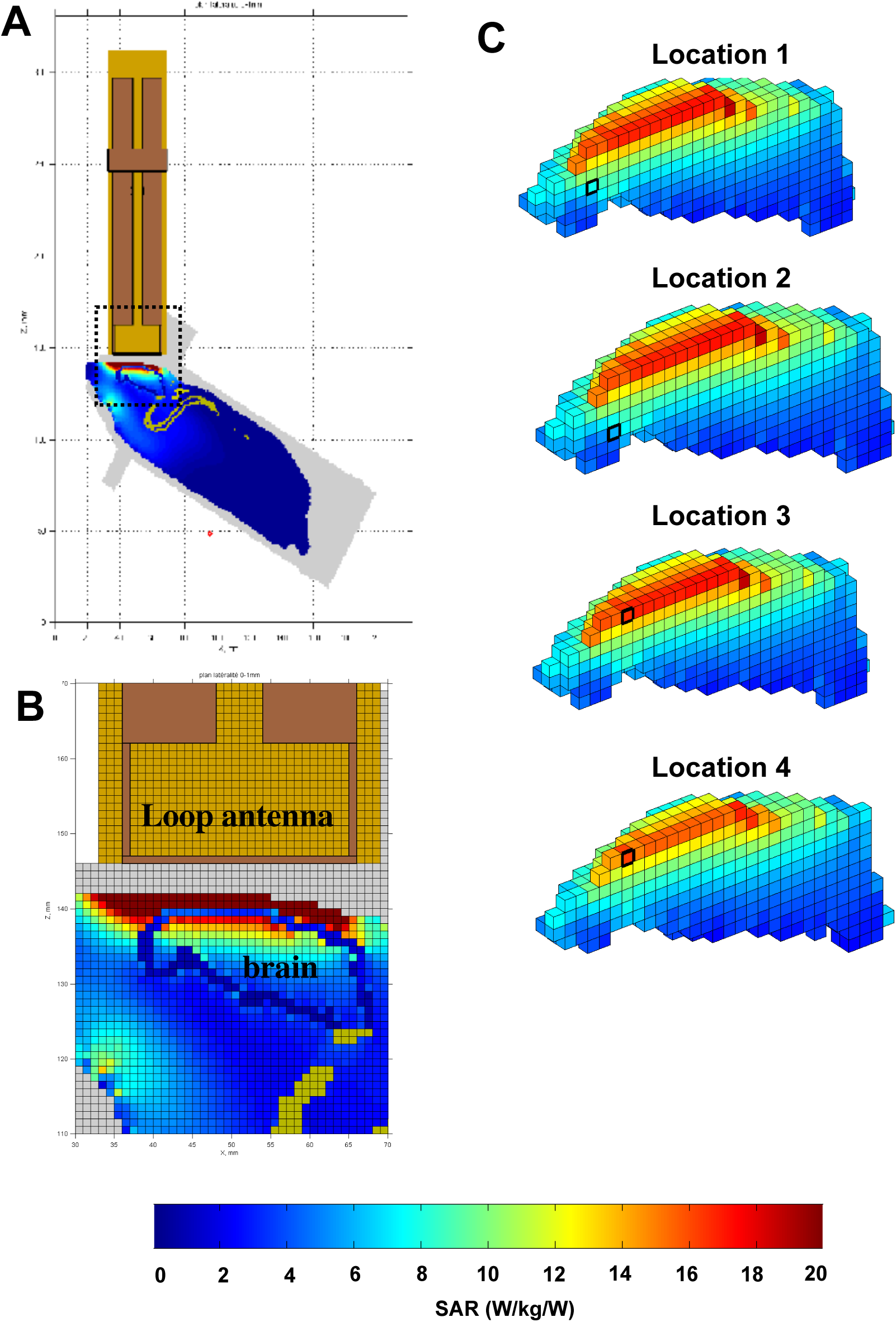
Dosimetric analysis of Specific Absorption Rates (SARs) in the rat brain during exposure to GSM 900MHz. The heterogeneous model of rat phantom and loop antenna described by Lévêque et al. (2004) was used to evaluate the local SAR in the brain with a 1 mm cubic mesh. As explained in Lévêque et al. (2004), MRI of a rat (250g) maintained in the rocket was carried out and the tissues (skin, skull, brain, marrow, fat, muscles) were segmented and assigned to different dielectric properties. A. Global view of the rat phantom in the rocket with the loop antenna above its head. The area defined by the dotted line is shown in B. B. This view shows the interface between the loop antenna and the rat’s head. It clearly indicates that the first millimetres of tissues below the loop antenna (i.e. the skin, the skull, the dura mater and the cortex) receive the highest SAR values. C. Based on the relative location of the brain areas in the Paxinos and Watson atlas (1986) and on their corresponding locations in the model of rat brain, values of SAR for the brain areas exhibiting changes in c-Fos labeling have been evaluated. The locations 1, 2, 3 and 4 show here correspond to the position of the prelimbic, infralimbic, frontal M2 and frontal M1 areas, respectively. Note that the two frontal areas (M1 and M2) receive high SAR values, whereas the prelimbic and infralimbic areas receive low SAR values because they are located deeper in the brain. Despite this important difference in terms of SAR values, they all expressed changes in c-Fos labeling.

## 4. Discussion

The aim of the present study was to evaluate, in a rat model, the potential health risks of exposure to 900MHz GSM signals. To mimic as closely as possible the use of cellular phones, we used a head-only exposure system allowing application of calibrated intensities of EMFs at the vicinity of the animal’s head. Mapping of brain regions affected by EMF exposure was achieved by measuring the expression of the c-Fos immediate early gene, which is rapidly induced after external stimulation. As c-Fos expression and behavioral performance can both be affected by stressful conditions (Sharp et al., 1991; Armario et al 2004), special care was taken to minimize as much as possible the stress effect induced by the confinement in exposure rockets. In the two experiments, we progressively habituated the animals to be restrained to reduce the stress. This procedure was probably successful because in experiment 2, no differences in c-Fos expression (in all but one brain area) and in behavioral performance were found between control-cage animals (animals maintained in their home cage) and sham animals (placed in the rockets but not exposed to EMFs). Although control-cage animals were not used in experiment 1, the extended habituation (two weeks vs. one week in experiment 2) should have largely reduced potential effects resulting from the restrained conditions.

In the first experiment, a 2h acute exposure to 900 MHz EMF affected differentially the brain regions analyzed in animals at rest. Out of the 23 brain areas investigated only six showed significant changes in c-Fos labeling with a dose-dependent increase for five of them at a BASAR of 1W/kg. In the second experiment, after 10 or 14 days of 45-min exposure to 900MHz EMF delivered immediately before behavioral testing, six brains areas displayed a significant decrease in c-Fos labeling. It is quite striking that four of these brain areas were cortical areas exhibiting increases in c-Fos labeling after a 2h acute exposure. Surprisingly, our dosimetric analysis indicated that among the brain areas showing changes in c-Fos labeling, some areas received high SAR values whereas others corresponded to low SAR values.

### 4.1. Relation with previous studies evaluating the effects of acute exposure to EMFs

Only a few studies have addressed the possibility that mobile phone radiations could induce genomic responses *in vivo*. Using head-mainly exposures, a slight increased expression of c-Fos messenger RNA was observed by Fritze et al. (1997) in some brain areas (cerebellum, neocortex and piriform cortex) after 4h exposures to various BASARs (0.3, 1.5 and 7.5 W/kg). As these increases were also detected on sham-exposed animals, they most likely result from the restrained conditions used during EMF exposure (see Finnie 2005). Furthermore, hyperthermia that has been measured in most rats following whole-body exposure to SAR > 5W/kg could explain the prominent c-Fos protein expression observed in several cortical and periventricular areas (Mickley et al., 1994). Different upstream regulatory elements could mediate an increase in c-Fos expression: changes in neurotransmitters releases, increase of Ca^++^ influx, neurotrophic factors. Also, emerging evidence suggests that c-Fos is essential in regulating neuronal cell survival versus death (Zhang et al., 2002). Previous *in vitro* studies have reported that different microwave exposures (continuous or pulsed) influence signal transduction pathways in brain tissue (Blackman et al., 1980; Gandhi & Ross, 1989). However, these experimental models often used EMFs quite different from the GSM signals and, therefore, the results should be interpreted with caution.

As the c-Fos immediate early gene is not a marker of cerebral injury, changes in c-Fos expression observed following EMF exposure do not necessarily indicate a deleterious effect of EMFs on brain metabolism. In a few recent *in vivo* studies using a head-only exposure system similar to ours, molecular and cellular alterations of brain cells have been described after acute exposures to 900MHz GMS signals. First, a 2h exposure at BASARs of 4 and 32 W/kg decreased the content in gamma-aminobutyric acid (GABA) in the cerebellum (Mausset et al., 2001). Second, a significant reduction in the number of NMDA receptors at the post-synaptic level has been observed in cortex and striatum after a brief (15 minutes) exposure to 900MHz GSM signals (Mausset-Bonnefont et al., 2004). Also, increases in glial immunoreactivity have been reported in the cortex, striatum and cerebellum (Mausset-Bonnefont et al., 2004; Brillaud et al., 2007). However, these latter effects were obtained at quite high BASARs levels (32 W/kg and 6W/kg), out of the range of normal SAR delivered by cell phones. Some of these effects, particularly those on glial reactivity, were not replicated at lower BASAR values of 2-3 W/kg (Watilliaux et al 2011).

#### Relation with previous studies evaluating the effects of subchronic exposure to EMFs

After daily 45-min exposures to EMFs behavioral performance was not impaired in a working memory task, and the minor impairment observed in the reference memory task on a subset of animals was not confirmed on a larger size population (n=12 animal/group). These data replicate our original observations (Dubreuil et al., 2002), which have been corroborated by other studies (Sienkiewicz et al., 2000; Dubreuil et al., 2003; Cobb et al., 2004; Cassel et al., 2004). Exposing animals to behavioral tasks had two main advantages. First, it allows detection of possible decreases in c-Fos expression which would be otherwise difficult to observe in resting animals (because of floor effect). Second, it allows detection of changes in the capacity of certain brain regions to be activated in response to cognitive demand. Regarding the first advantage, we observed that the level of c-Fos expression in the sham-exposed rats differed in the acute versus the subchronic exposure experiment. For example, the mean value obtained for the prelimbic area was about 310 in experiment 1 (acute exposure), whereas it was 495 and 560 (in the working memory and in the reference memory task, respectively) in experiment 2 (subchronic exposure). These large increases (60% and 80%) in c-Fos labeling in sham animals probably indicate the involvement of several brain areas in the behavioral tasks during experiment 2.

Two hypotheses could be proposed to account for the decrease in c-Fos labeling observed in exposed animals during the second experiment. First, one could consider that neurons, as any biological tissue (Di Carlo et al., 2000, 2001), adapted to repeated exposure to EMF, and responded differently to an acute vs. a subchronic exposure. Thus, although the behavioral tasks increased c-Fos expression in all animals, this increase was attenuated in the animals daily exposed to the EMF because the repetition of exposures produced opposite effects when compared with a single exposure. However, according to this hypothesis, we should have obtained the same decreases after the two behavioral tasks, which was not the case. A second explanation relies on the fact that performing a behavioral task produced a “cognitive demand” which led to activation of several cortical networks, an activation which was attenuated by prior exposure to the GSM EMFs. This possibility may explain why the results differed between the two behavioral tasks: In the easiest task, the reference memory task, EMF exposure-induced decreases were limited to temporal and visual cortices. In contrast, in the working memory task, animals exposed to EMFs exhibited a reduction in the number of c-Fos-positive nuclei in many cortical regions (prelimbic, infralimbic, frontal, cingulated, perirhinal, and visual cortices; area CA1 in hippocampus) suggesting the recruitment of a larger network of brain areas during this task.

As the c-Fos protein is widely considered as an indirect correlate of neuronal activation (Sagar et al., 1988), the present data point out that daily exposure to GSM signals attenuates the increase in neuronal activity normally observed in brain areas during a behavioral task. However, this hypo-activity was not sufficient to induce detectable concomitant deficit performance in the working and reference memory tasks used here. One may argue that the use of more challenging behavioral tests might have revealed cognitive impairment following EMF exposure. However, we could not found impairments either by using a more complex working memory task (introducing a 10s confinement between the visit of each arm or a 15-min delay after 4 visited arms) or by increasing the temporal delay between the acquisition and the testing period in an object recognition task (Dubreuil et al., 2003). Several additional experiments would be necessary to discard totally the possibility that the decrease in c-Fos labeling detected here does not lead to behavioral deficits. For example, the use of longer retention interval associated with a reference memory task (e.g. retention sessions given several weeks after the end of acquisition) could reveal detectable retrieval deficits. It is also possible that longer subchronic exposures (over weeks) might reveal behavioral impairments with the same paradigms used here.

Recently, cellular alterations of brain cells have been reported after subchronic exposures to 900MHz GSM signals. A decrease in cytochrome oxydase activity was detected in several cortical areas (including the frontal areas) after a 7 days exposure to BASARs of 6W/kg but not after a similar duration of exposure to BASARs of 1.5W/kg (Ammari et al., 2008b). Longer exposures (5 days a week for 24 weeks) also induced increases in GFAP immunostaining in several brain areas (frontal cortex, dentate gyrus, striatum) at a BASAR of 6W/kg but not at a BASAR of 1.5 W/kg (Brillaud et al., 2007; Ammari et al., 2008a). Using low BASAR levels (1.5 and 3.5W/kg), the present data show that exposure to EMFs do have biological effect on neurons but also indicate that changes in neuronal activity (as revealed by c-Fos imaging) do not necessarily lead to behavioral impairments. As cytochrome oxydase activity is an endogenous metabolic marker, the effects that it can reveal (as well as those revealed by GFAP immunostaining) might reflect more permanent changes obtained when the neural tissue is chronically exposed to higher SARs (6W/kg in the case of the study by Ammari et al 2008).

#### Open questions and conclusions

Several questions remain open after the present study. The first and more puzzling one is the dose-dependent increase in c-Fos labeling observed in experiment 1: in 5/6 brain areas, the BASAR level which produces an increase in c-Fos labeling was not the highest BASAR tested (6W/kg) but the BASAR of 1W/kg. The existence of windows for the effects of EMFs (either for the frequency, the duration or the intensity of the EMF) has long been debated and the cellular bases of these effects are still unclear (see for review Adey, 1993; Adair, 1997; Berg, 1999). In the range of SAR used here, it was shown that the incidence of DMBA-induced mammary gland tumors was increased by subchronic exposure to 900MHz GSM at a SAR of 1.4W/kg but not at 2.2 or 3.5W/kg (Anane et al., 2003). In vitro studies indicate (i) that neurons are at least 10 times more sensitive than previously thought to weak electric fields (Francis et al., 2003), (ii) that electrophysiological effects on neuronal activity can be obtained at very low SARs (0.0016-0.0044 W/kg) and (iii) that within this range the effects can differ as a function of the field intensity (Tattersall et al., 2001). The second unresolved question is why some brain areas expressed changes in c-Fos labeling whereas others did not. The dosimetric analysis allows discarding the possibility that changes in c-Fos labeling only occur in brain areas receiving the highest SARs. However, it is possible that unspecific stimuli, such as local thermal stimuli occurring during exposure within the skin or the dura, activated brain regions even if these brain regions did not receive the highest SARs. This possibility should be kept in mind because many studies in human have shown that noxious or innocuous thermal stimuli activate cortical areas such as the anterior cingulate cortex and other regions of the frontal cortex (Kwan et al., 2000, Moulton et al., 2005; Seifert et al., 2007, Gundel et al., 2008) which are similar to those showing changes in c-Fos labeling in our experiments. In fact, a recent study showed that very focal EMF exposure of the cortical tissue can produce an increase in brain temperature and in local cerebral flow (Masuda et al 2011). Last, it is also possible that some brain areas are more sensitive to EMF than others. In this context, it is important to mention that, in human, a 30 minutes exposure to 900MHz GSM signals (BASAR 1W/kg) has been shown to increase the relative regional cerebral blood flow in the dorsolateral prefrontal cortex on the side of exposure (Huber et al., 2005). By showing that significant effects can be obtained at low SAR levels, the present results obviously call for more systematic investigations looking for changes in neuronal activity after long duration exposures to low SAR levels.

## Acknowledgments

The authors have no conflict of interest to declare.

The present work was supported by the COMOBIO project carried out within the RTRN research program (National Network for Research in Telecommunication). DD was supported by France Telecom R&D. Our funding sources had no involvement in the collection, analysis and interpretation of the data, in the writing of the manuscript and in the decision to submit the paper for publication.

We thank Dr. Rachid Anane (PIOM laboratory, now IMS laboratory, Bordeaux) for his help during some experiments performed in Bordeaux.

## List of Abbreviations

BASAR: Brain Average Specific Absorption Rate
EMF: Electromagnetic Field
GSM: Global System for Mobile communication
RM: Reference Memory
SAR: Specific Absorption Rate
WM: Working Memory

---

**Supplementary Figure 1:**
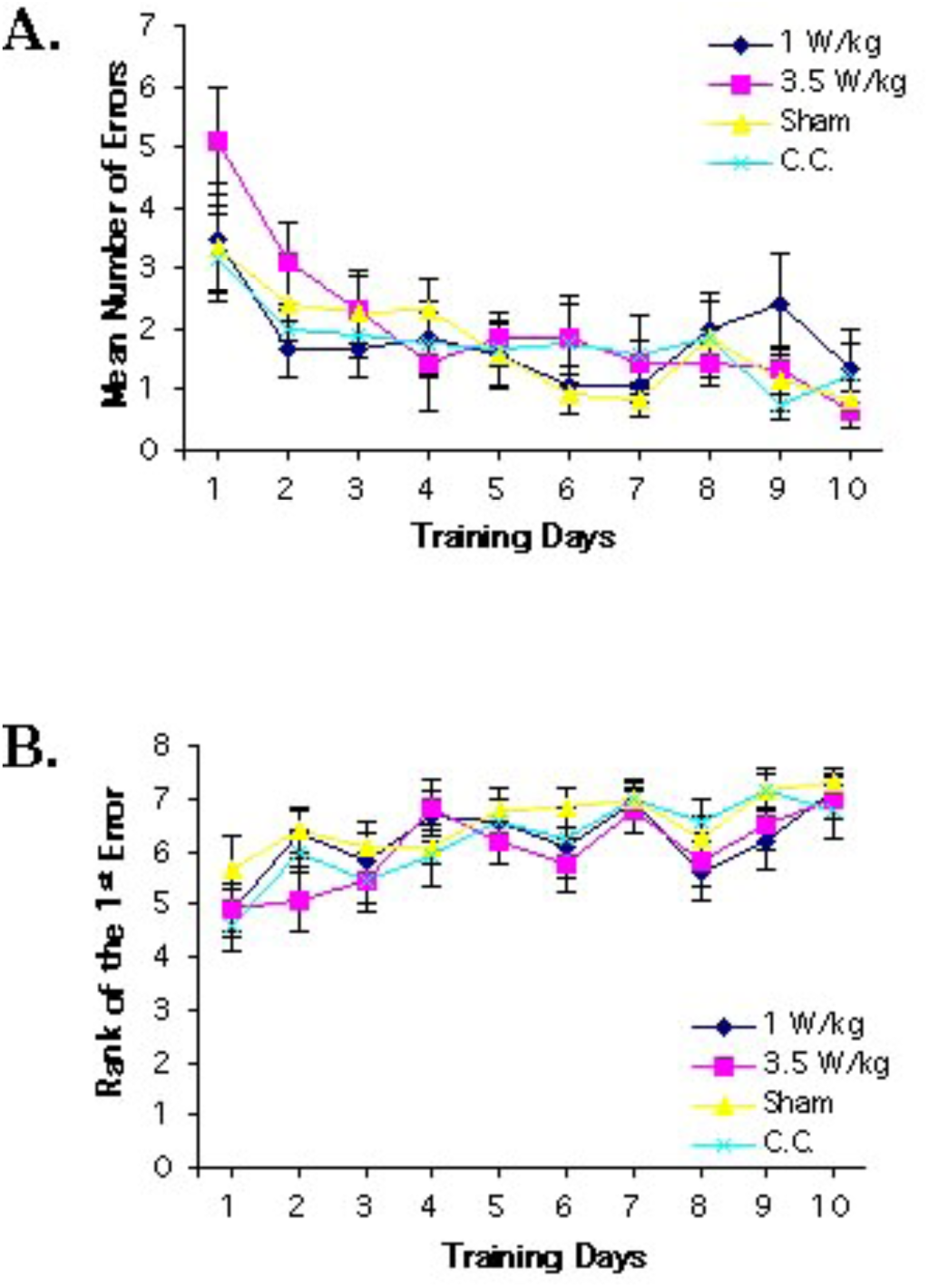
Performance in radial-arm maze (working memory task) for the total popula?on of animals (n=12/group): (A) The mean number of errors decreased from day to day for all the treatment groups. There was no significant difference between the groups. (B) The rank of the first error increased from day to day for all groups. The learning curves of exposed and non-exposed rats are similar. As assessed by the number of errors and the rank of the first error, all groups learned in a similar way.

**Supplementary Figure 2:**
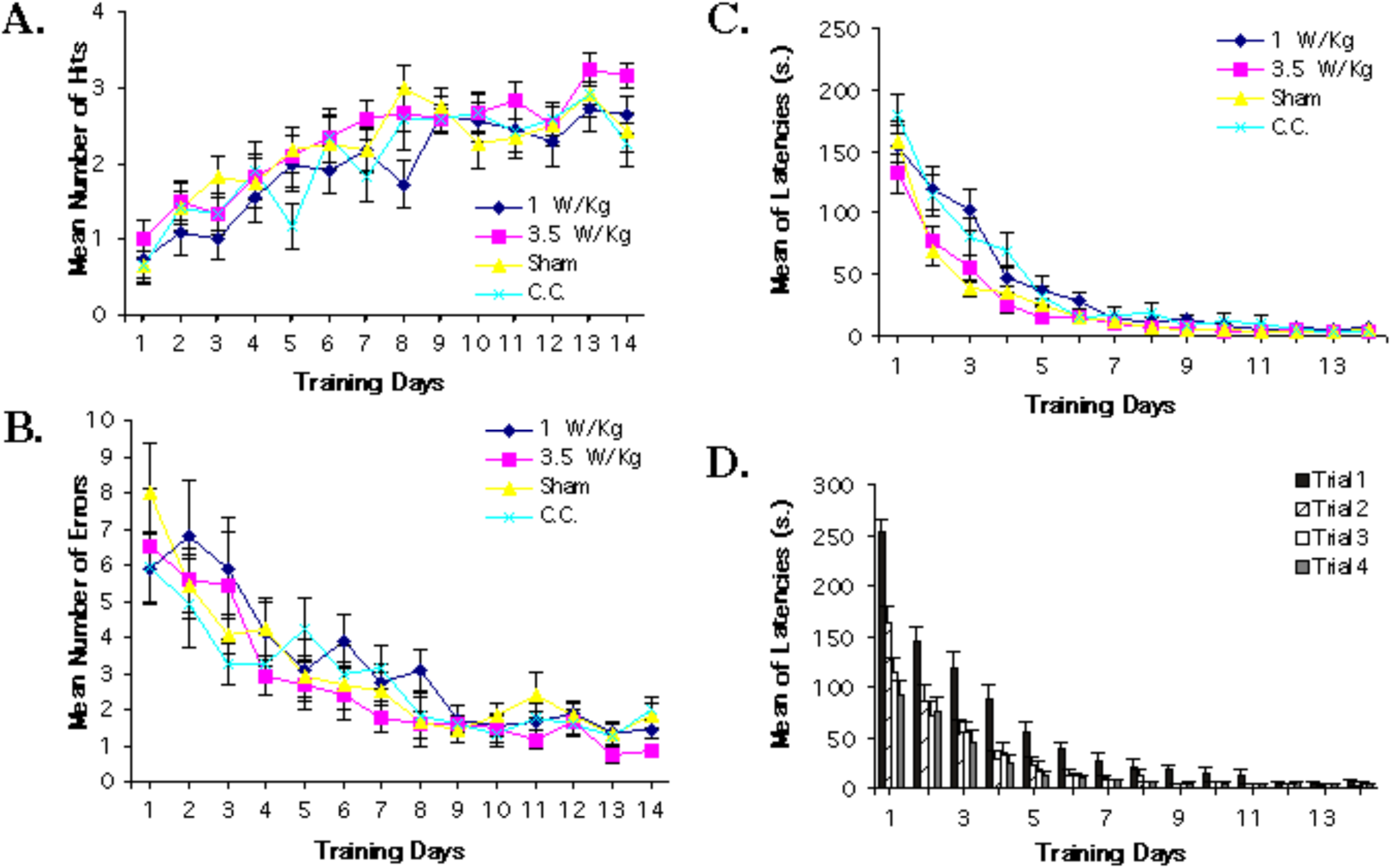
Performance of the whole popula?on of rats (n=12/group in the reference memoy task. (A) The number of hits increased over the whole training period for all the groups. There was no significant difference between the learning performance of the treatment groups. (B) The number of errors decreased during the training period for all groups. There was no significant difference between the learning performances of the treatment groups. (C) The latencies decreased similarly for all treatment groups. (D) The latency of the first trial was always longer than the other trial latencies (warm-up decrement effect). The difference between the latency of the first trial and the other latencies diminished over the days and this aOenua?on was similar for the treatment groups.

**Supplementary table 1.**

|  | Anteriority | Laterality | Depth | SAR W/kg/W |  |
| --- | --- | --- | --- | --- | --- |
| PrL Area | 4-5 mm | 0-1 mm | 3-4 mm | 7.80 | Point 1 |
| IL Area | 4-5 mm | 0-1 mm | 4-5 mm | 5.74 | Point 2 |
| FM2 Area | 5,7-6.7 mm | 0-1 mm | 0-1 mm | 16.44 | Point 3 |
| FM1 Area | 5.7-6.7 mm | 2-3 mm | 0-1 mm | 16.06 | Point 4 |
| Dorsal Septum | 5.7-6.7 mm | 0-1 mm | 3.5-4.5 mm | 8.90 | Point 5 |
| Ventral Septum | 5.7-6.7 mm | 0-1 mm | 5-6 mm | 5.75 | Point 6 |
| Cg2 Area | 6.7-7.7 mm | 0-1 mm | 2-3 mm | 10.58 | Point 7 |
| Parietal Area | 9.5-10.5 mm | 5-6 mm | 0-1 mm* | 8.72 | Point 8 |
| Subiculum | 12-13 mm | 1-3 mm | 3-4 mm | 8.37 | Point 9 |
| Visual Area | 12-13 mm | 5-6 mm | 0-1 mm* | 7.72 | Point 10 |
| Temporal Area | 12-13 mm | 6.5-7.5 mm | 0-1 mm* | 7.720 | Point 11 |

## Footnotes

1 Initially, we aimed at a BASAR of 4W/kg to match the exposure of one group of experiment 1; but we could not reach this value with our exposure system.

